# Global distribution of a land plant by means of oceanic dispersal

**DOI:** 10.1101/2024.12.28.629971

**Authors:** Mohammad Vatanparast, Koji Takayama, Yoichi Tateishi, Tadashi Kajita

## Abstract

The geographic range of species is one of the key indicators of their evolutionary success. Unlike animals, most plants are immobile and expand their range primarily through long-distance dispersal (LDD) of seeds. Seeds can be dispersed over very long distances, especially by ocean currents. However, little is understood about the role of ocean currents in LDD and the contribution of land barriers to genetic differentiation on a global scale. To investigate the outermost limits of LDD by ocean currents in land plants, we studied *Canavalia rosea* (Sw.) DC. (Fabaceae), which is distributed throughout littoral areas of the tropics and subtropics around the world. Our results using DNA sequences from six chloroplast regions and two low-copy nuclear loci from 436 individuals in 37 populations revealed that ocean currents played a crucial role in the species’ range expansion and maintaining its global distribution despite the constraints by continental barriers. These suggest that the global distribution of *C. rosea* is the result of recent transoceanic dispersal within and between the Atlantic and Indo-Pacific oceans, while the Isthmus of Panama has prevented gene flow across the American continents. These results highlight the power of oceanic seed dispersal in shaping plant biogeography.

---

The geographic range is a key indicator of the evolutionary success of living organisms^1,2^. Unlike animals, plants are immobile and have adapted long-distance dispersal (LDD) of seeds and propagules to expand their geographic range^3^. The major barriers to plants using LDD are the oceanic gaps between continents and the continents themselves. Many biogeographic studies are invoking LDD over ocean or continental barriers in shaping global distributions of plants^4–6^. However, little is understood about the role of ocean currents in LDD and the contribution of land barriers to genetic differentiation on a global scale^7^. Pantropical plants with drifting seeds provide extreme models for studying these patterns^8,9^. The beach bean, *Canavalia rosea* (Sw.) DC. (Fabaceae) is a pantropical leguminous vine that grows on sandy beaches in tropical and subtropical regions. Its distribution range is consistent with the areas where the average surface ocean water temperature is around 20°C. Seeds of *C. rosea* can float in seawater for months while remaining impermeable^10^, which enables this species to achieve one of the widest global distributions among pantropical plants with drifting seeds^11^. We sampled 436 individuals from 37 populations covering the entire distribution range and analyzed DNA sequences of six chloroplast regions (cpDNA) and, for a subset of individuals, two low-copy nuclear loci (*LEAFY* and *RNAH*) to assess transoceanic seed dispersal patterns.

Our results revealed 17 cpDNA haplotypes, with the population genetic structure shaped by the American and African continents (Fig. 1). The H8 (orange) and H6 (green) haplotypes showed very broad distributions in the Indo-Pacific and Indo-Atlantic oceans, respectively. While the distribution of haplotypes on the same continent could result from gradual range expansion, the existence of common haplotypes across oceanic regions strongly suggests LDD by ocean currents given the recent history of the species (<5 Mya^12^). The haplotype H8 is present in the Indian Ocean and across the Pacific Ocean, a region that typically acts as a major barrier for sea-dispersed plants like mangroves^13^ and *Ipomoea pes-caprae* (L.) R.Br. (Convolvulaceae)^14^. The nuclear loci haplotype distributions are in agreement with cpDNA haplotype results (Fig. S1, Fig. S2).

**Figure 1:**
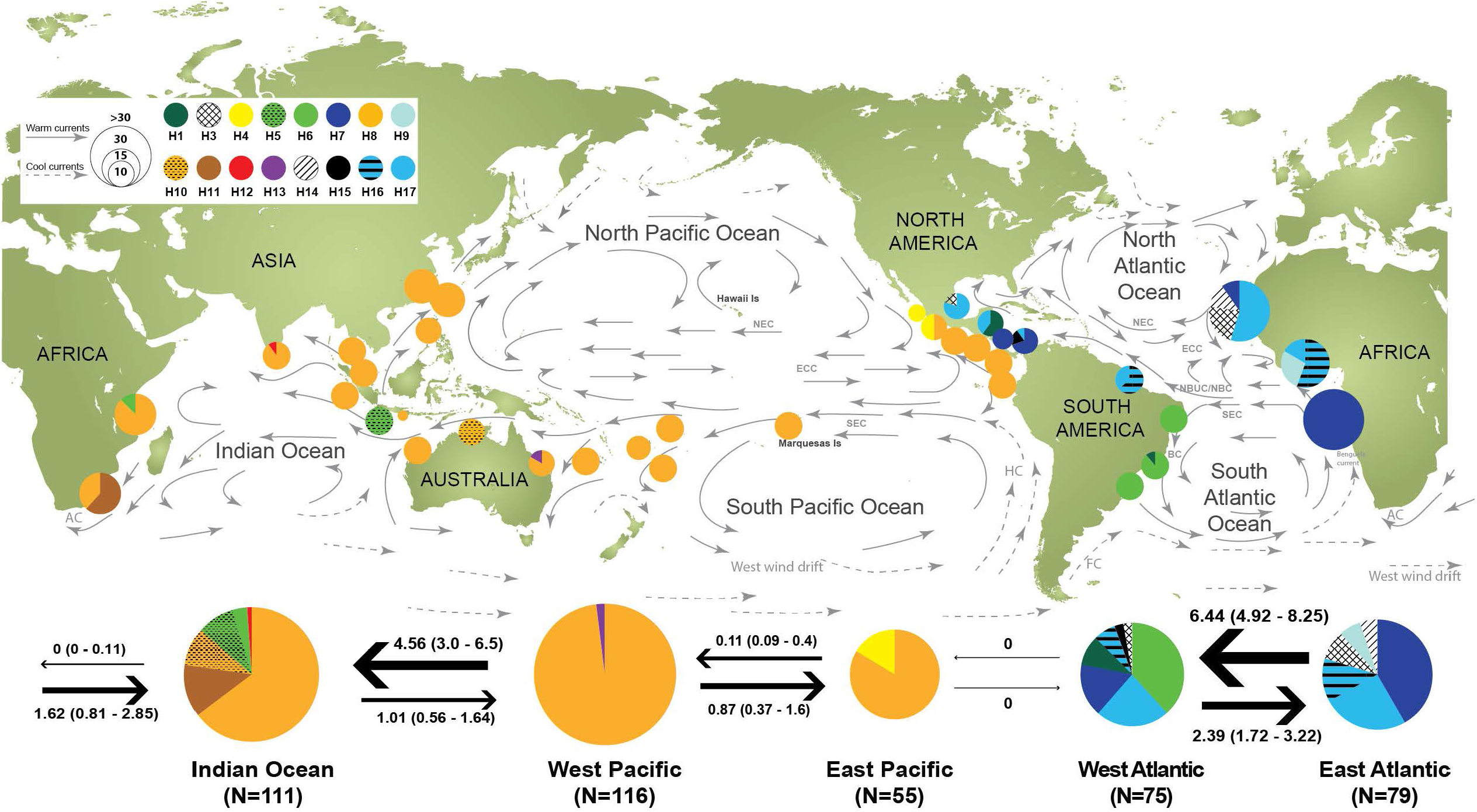
Distribution of the chloroplast haplotypes of *C. rosea* based on partial sequence data (2,012 bp) of six cpDNA regions. The haplotypes (H7, H19-21) and (H16, H18) in the full-length data (Fig. 2) were collapsed into a single haplotype H7 and H16, respectively. Pie charts represent the proportion of haplotypes at each locality, and the size of the pie charts is proportional to the sample size. Black arrows indicate the direction of migration and the number of migrants (*N*_*m*_) estimated using the MIGRATE-N for five regional groups (West Atlantic, East Atlantic, Indian Ocean, West Pacific, and East Pacific). Ranges in parentheses show the minimum and maximum values derived from three independent runs. Surface ocean currents (warm and cool) are depicted with gray arrows (redrawn from^25^): AC: Agulhas Current, BC: Brazil Current, ECC: Equatorial Countercurrent, FC: Falkland Current, HC: Humboldt Current, NBUC/NBC: North Brazil Undercurrent/Current, NEC: North Equatorial Current, SEC: South Equatorial Current.

The geographic distribution of cpDNA and nuclear haplotypes (Fig. 1, Fig. S1, Fig. S2) and the cpDNA haplotype network (Fig. 2) show likely paths of range expansion by ocean currents. The cpDNA haplotype network identifies haplotype H7 from the Atlantic as ancestral, based on statistical parsimony analysis. Other haplotypes diverged from H7 to form distinct geographic clusters. In particular, Atlantic haplotypes diversified by a few mutations into Indo-Pacific oceans haplotypes (H4, H8, H12, H13). The Pacific haplotype H8, in turn, connects to Western Atlantic haplotypes (H5, H6) independently of other Atlantic haplotypes, suggesting bidirectional exchange via the Indian Ocean. The higher haplotype diversity observed in the Atlantic region (Fig. 1, Table S1) supports an Atlantic ancestry hypothesis. In the network results, the Hawaiian endemic *Canavalia* DC. species are related to the Atlantic H7, suggesting distribution in both regions prior to the closure of the Isthmus of Panama (Fig. 2). Molecular clock analyses indicate that the Hawaiian endemic species clade is less than 5 million years old^12^, suggesting that these species recently diversified in the Pacific. Statistical inferences reveal strong genetic structure across the Panama Isthmus (Table S2), with no migration (*N*_*m*_ = 0) between eastern Pacific and western Atlantic populations (Fig. S3), similar to patterns observed in *Hibiscus tiliaceus* L. (Malvaceae)^8^ and seabirds^15^. These results emphasize the role of the Panama Isthmus closure as a barrier to ocean current-mediated plant dispersal.

**Figure 2:**
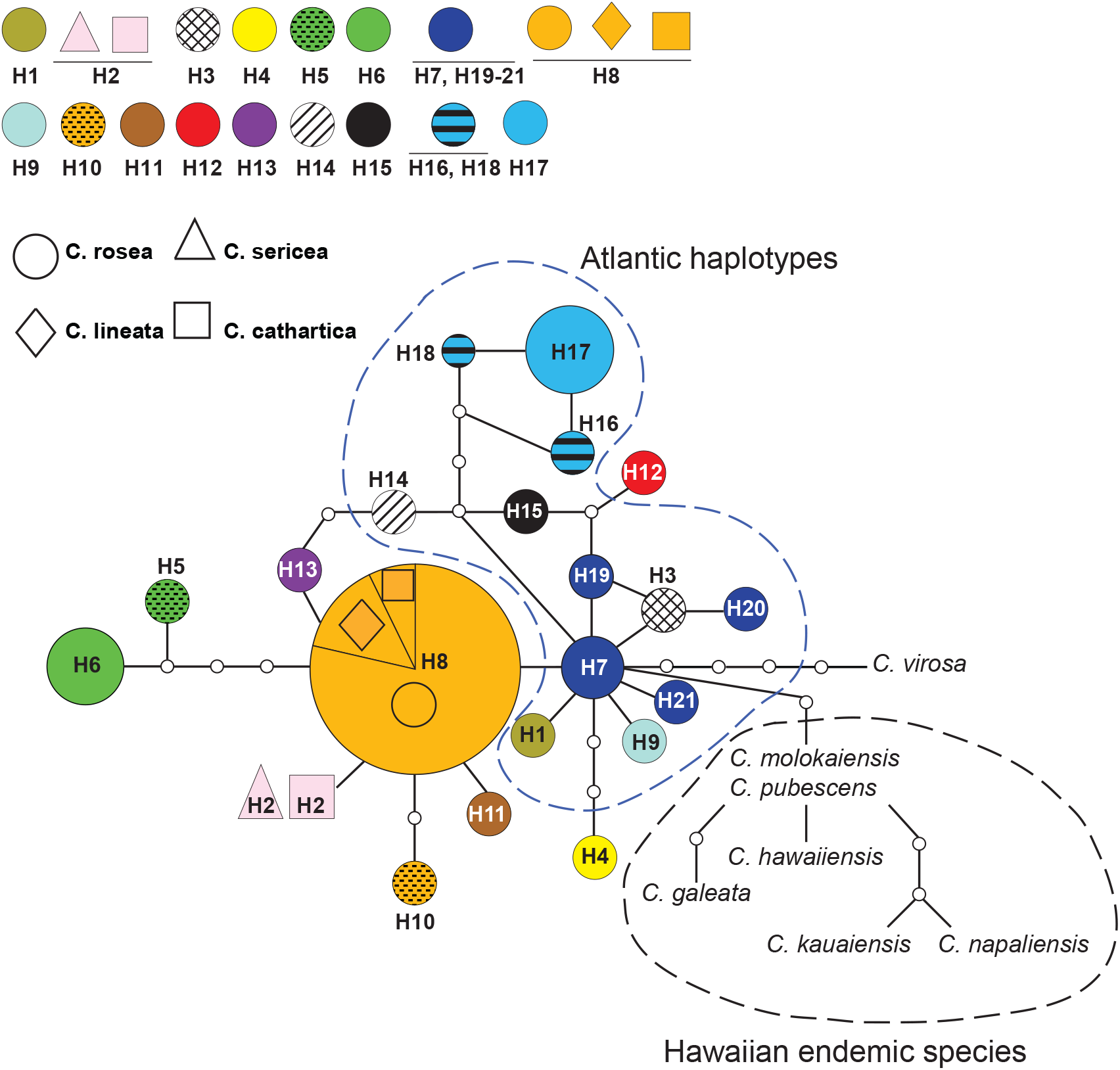
Statistical parsimony networks among *C. rosea* and its allied species based on the full length data of six cpDNA regions (ca. 6,400 bp; H1-21). Symbols, colors, and sizes of each haplotype correspond to the species, haplotypes, and frequencies of the corresponding haplotypes, respectively. The small empty circles indicate the undetected intermediate haplotypes.

Here, we demonstrate that LDD by ocean currents has expanded the geographic range of *C. rosea* along prevailing global trajectories, while land masses shaped distinct evolutionary lineages. Using global sampling and molecular markers, we found that the American continents prevent seed dispersal between the Pacific and Atlantic coasts, while migration occurs across the other side of the globe. This creates a pantropical distribution with genetic signatures similar to those of a ring species^16^. Our results show that LDD by ocean currents contributes to global plant distribution, while continental land barriers shape population structure and prohibit panmixis. Additional experiments are required to examine the population dynamics across the African and American continents.

## Summary of methods

Leaf samples were collected from the entire range of *C. rosea* for 436 individuals from 37 populations. Voucher specimens were deposited in the herbarium of the University of the Ryukyus (RYU) and in the Jardim Botânico, Rio de Janeiro (RBRJ). DNA extractions, PCR, and sequencing methods are described in Vatanparast *et al*. [9]. For the cpDNA haplotype distribution analysis (Fig. 1), we targeted partial sequences of six cpDNA regions based on our phylogenetic analyses^9^ and the final aligned sequences were 2,012 bp. For the cpDNA haplotype network analysis (Fig. 2), we used the full length of the six cpDNA regions (ca. 6,400 bp) for a subset of samples. The haplotypes (H7, H19-21) and (H16, H18) in the full length data (Fig. 2) were collapsed into a single haplotype H7 and H16 in partial sequences, respectively. For the nuclear markers, direct sequencing of the *LEAFY* and the *RNAH* loci were used in forward and reverse directions after the initial screening steps using primers from Choi *et al*. [17]. The number of individuals and alignment length for the *LEAFY* and the *RNAH* loci were 89/605bp and 60/746bp, respectively. The numbers of haplotypes (H), polymorphic site (S), Haplotype diversity (Hd), and nucleotide diversity (*π*) were calculated using the DnaSP version 5.10^18^. The genealogical relationships of haplotypes were constructed for the cpDNA, *LEAFY* and the *RNAH* data independently with the TCS v.1.23^19^ using the statistical parsimony method with a 95% confidence interval. To determine genetic differentiation among populations of *C. rosea* we carried out F-statistics (*F*_*ST*_) analysis of cpDNA data using the Arlequin v.3.5^20^. Pairwise *F*_*ST*_ was calculated following the method of Weir & Cockerham [21] and statistical significance was assessed using 1000 permutations. An exact test of population differentiation was performed to examine the null hypothesis of the random distribution of haplotypes across populations^22^. To assess migration rates, the direction of gene flow, and genetic diversity among populations, the Maximum Likelihood and Bayesian approaches implemented in MIGRATE-N v.3.1.3^23^ were used. We were specifically interested in contrasting the pairwise migration rate (M) and direction of gene flow between geographic regions rather than between sampled populations. Hence, all populations were pooled into five sub-oceanic regions (Indian Ocean, West and East of each Atlantic and Pacific Oceans). Given the linear distribution of *C. rosea* at coastal lines, the stepping stone migration model with asymmetric rates^24^ was employed in MIGRATE-N.

## Acknowledgments

This study was supported by grants from the Japan Society for the Promotion of Science KAKENHI 15K14588, 19370032, 16370043, and 14740471 to T. K.; 14405015 and 12575011 to Y. T. and a grant from the Environment Research and Technology Development Fund (S-9). We thank H. C. Lima, M. Bovini, A. Tidiane Ba, D. Darnaedi, I. Ibrahim, D. Neil, D. A. Madulid, E. Narvaez, I. Seneviratne, J. Flora, T. Yahara, A. Soejima, T. Ohi-Toma, K. Watanabe, Y. Mori, Y. Kita, Y. Abe, BISH (USA), JBRJ (Brazil), CNPq (Brazil), STRI (Panama), OTS (Costa Rica) and MEXU (Mexico) for their support in field sampling. We also thank Y. Watano and T. Asakawa, for their useful comments. Our special thanks to R. Toby Pennington and Ashley N. Egan for reading the early versions of the manuscript. M. V. was supported by a PhD scholarship from Japan’s Ministry of Education, Culture, Sports, Science and Technology (MEXT).

## Data Availability

The datasets generated in the current study are available from the corresponding authors.

---

**Table S1:** Number of individuals (N), Number of haplotypes (h), Polymorphic sites (S), Haplotype diversity (Hd), and Nucleotide diversity (*π*) across five sub-oceanic regions of *C. rosea*.

| Regions | N | (h) | (S) | (Hd) | ($\pi$) |
| --- | --- | --- | --- | --- | --- |
| Indian Ocean | 111 | 6 | 8 | 0.5507 | 0.00053 |
| West Pacific | 116 | 2 | 1 | 0.0341 | 0.00002 |
| East Pacific | 55 | 2 | 3 | 0.2787 | 0.00042 |
| West Atlantic | 75 | 7 | 8 | 0.7809 | 0.0014 |
| East Atlantic | 79 | 6 | 6 | 0.7403 | 0.00087 |

**Table S2:**
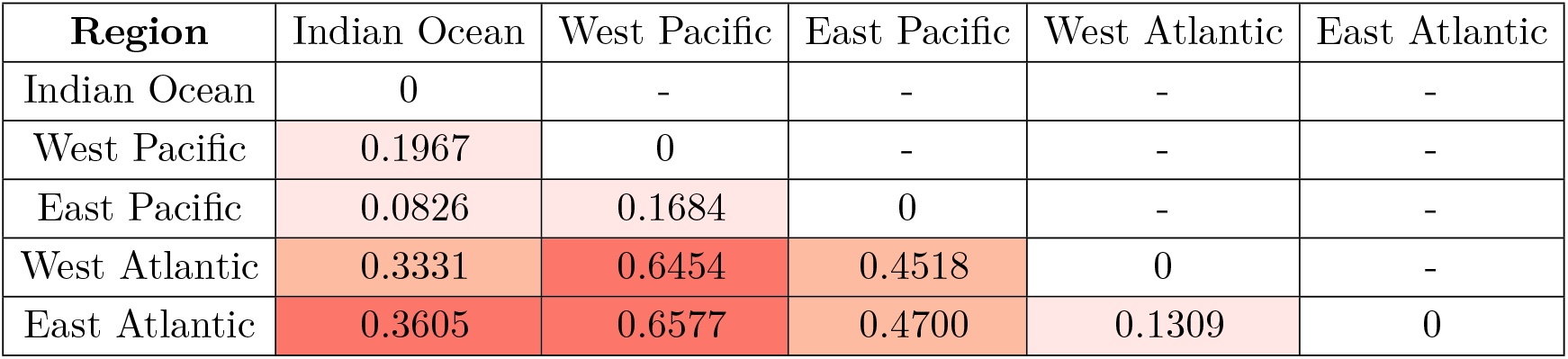
Pairwise *F*_*ST*_ comparison between five sub-oceanic regions of *C. rosea*. All differentiation results are significant based on the exact test (*P <* 0.05).

**Figure S1:**
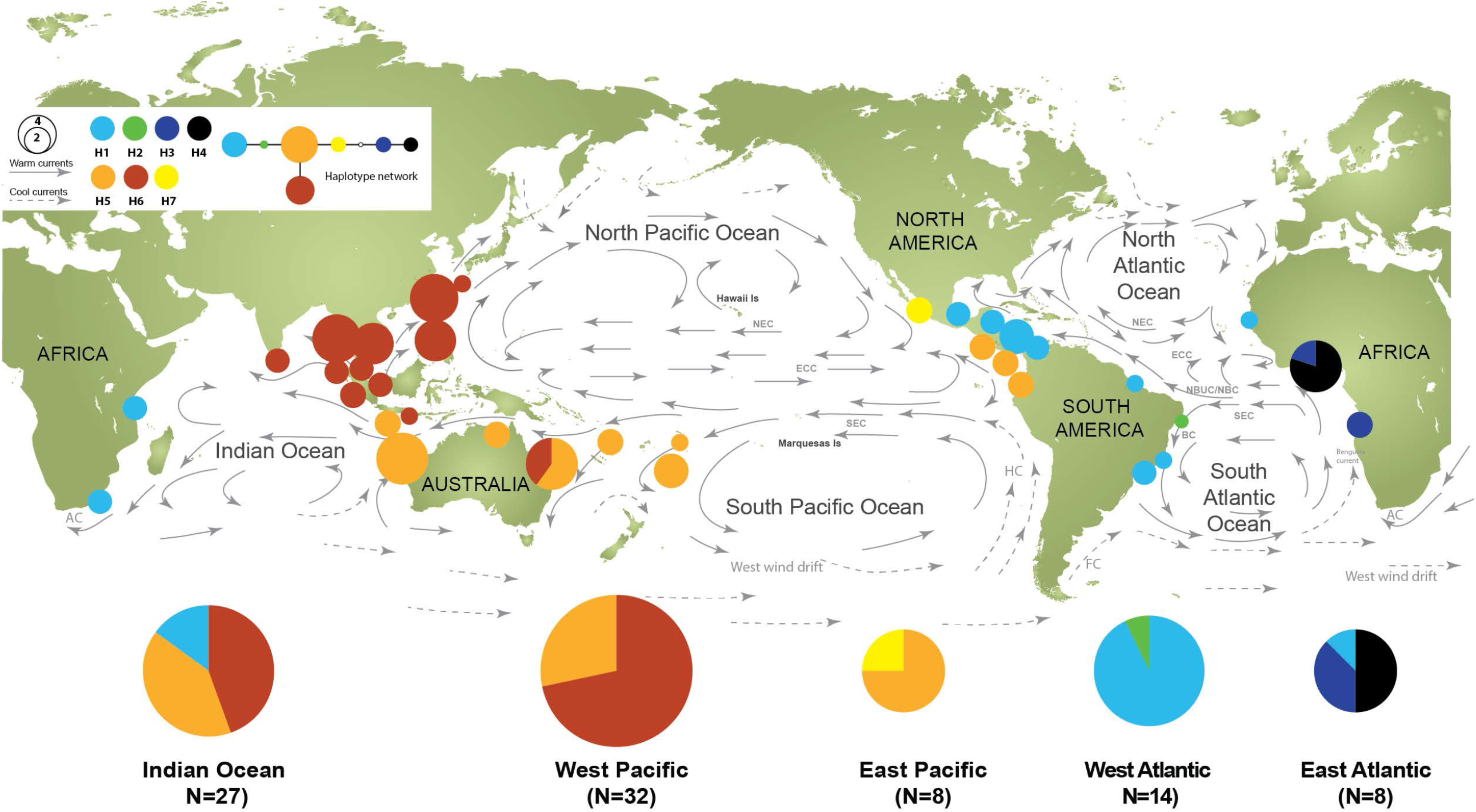
Distribution of the nuclear *LEAFY* haplotypes of *C. rosea*. See (Fig. 1) for the details.

**Figure S2:**
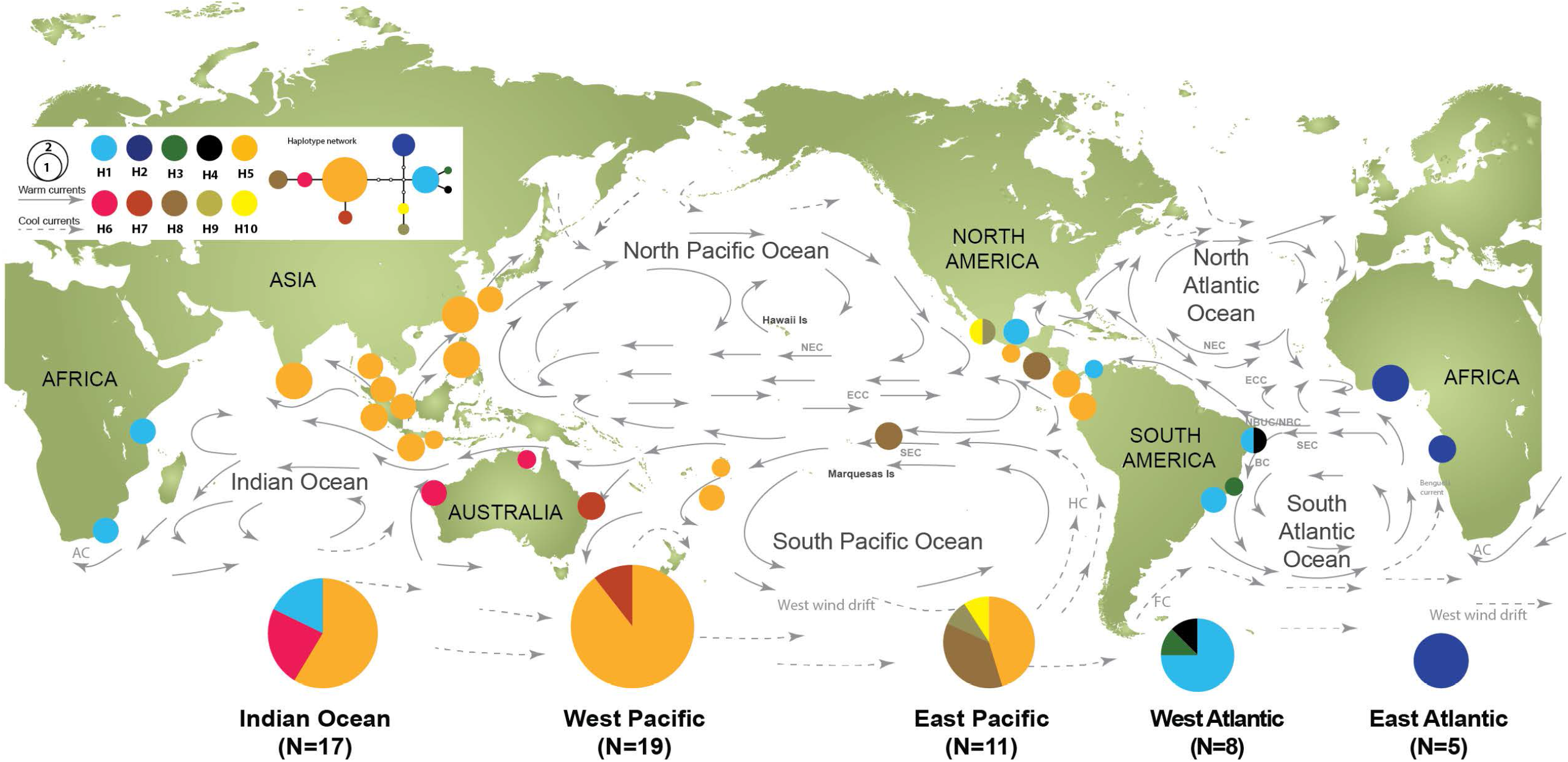
Distribution of the nuclear *RNAH* haplotypes of *C. rosea*. See (Fig. 1) for the details.

**Figure S3:**
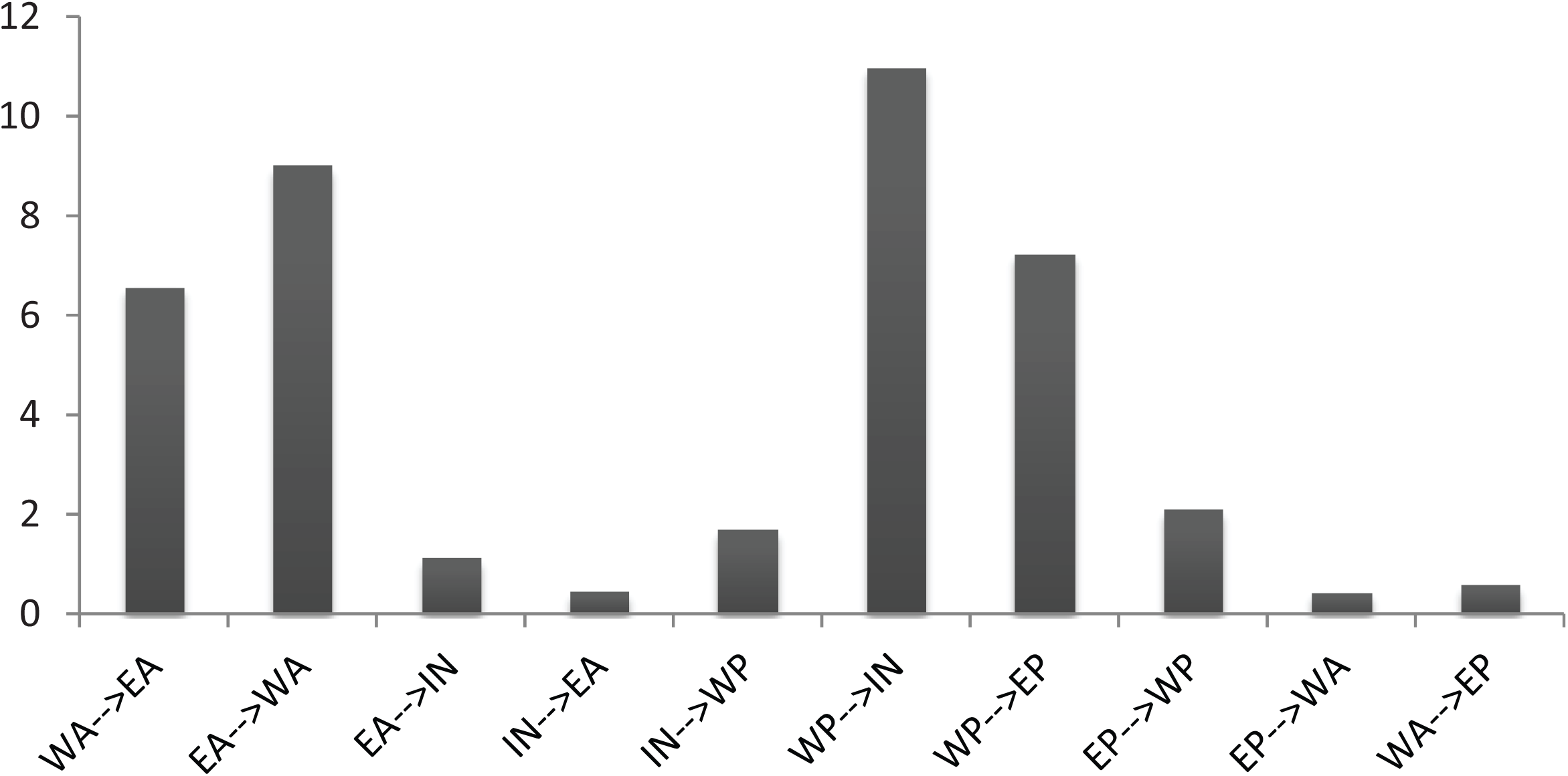
Estimated number of migrants (*N*_*m*_) between five sub-oceanic regions obtained from Bayesian method implemented in MIGRATE-N^26^. The calculation was based on a 95% credible interval of *θ* and M values. WA: West Atlantic, EA: East Atlantic, In: Indian Ocean, WP: West Pacific, EP: East Pacific.

